# When metabolism meets physiology: Harvey and Harvetta

**DOI:** 10.1101/255885

**Authors:** Ines Thiele, Swagatika Sahoo, Almut Heinken, Laurent Heirendt, Maike K. Aurich, Alberto Noronha, Ronan M.T. Fleming

**Affiliations:** Luxembourg Centre for Systems Biomedicine, University of Luxembourg, Campus Belval, Esch-sur-Alzette, Luxembourg; Faculty of Science, Technology and Communication, University of Luxembourg, Campus Belval, Esch-sur-Alzette, Luxembourg.; Division of Analytical Biosciences, Leiden Academic Centre for Drug Research, Faculty of Science, University of Leiden, Leiden, The Netherlands.

## Abstract

Precision medicine is an emerging paradigm that requires realistic, mechanistic models capturing the complexity of the human body. We present two comprehensive molecular to physiological-level, gender-specific whole-body metabolism (WBM) reconstructions, named Harvey, in recognition of William Harvey, and Harvetta. These validated, knowledge-based WBM reconstructions capture the metabolism of 20 organs, six sex organs, six blood cells, the gastrointestinal lumen, systemic blood circulation, and the blood-brain barrier. They represent 99% of the human body weight, when excluding the weight of the skeleton. Harvey and Harvetta can be parameterized based on physiological, dietary, and omics data. They correctly predict inter-organ metabolic cycles, basal metabolic rates, and energy use. We demonstrate the integration of microbiome data thereby allowing the assessment of individual-specific, organ-level modulation of host metabolism by the gut microbiota. The WBM reconstructions and the individual organ reconstructions are available under <u>http://vmh.life</u>. Harvey and Harvetta represent a pivotal step towards virtual physiological humans.

## Introduction

At the heart of precision medicine lies a predictive model of the human body that can be interrogated in a personalized manner for potential therapeutic approaches *in-silico (1).* While molecular biology has yielded great insight into the ‘parts list’ for human cells; so far, limited progress has been made for integrating these parts into a virtual whole human body. The Virtual Human Physiome project has generated comprehensive knowledge about the working of the human body and its organs (*2*) but has yet to be connected with molecular-level processes and their underlying networks of genes, proteins, and biochemical reactions.

Although an *in-silico* molecular-level description of humans metabolism is available (*3*), the generation of accurate organ-and tissue-specific metabolic models remains challenging using automated approaches and omics data (*4*). At the same time, a solely manual curation approach based on extensive literature review is not tractable due to the large number of organs and cell-types in the human body as well as the fact that organs and their metabolic functions have been studied at a different depth. Hence, a combined algorithmic and manual curation approach is needed, which has already been applied to microbial metabolic models (*5*).

To contribute towards the ambitious goal of a whole-body model of human (*6*), current systems biology approaches need to go beyond human metabolism, by accurately including anatomical and physiological properties in the computational modeling framework, such as constraint-based modeling (*7*) and flux balance analysis (*8*). For instance, Bordbar et al. connected three organ-specific metabolic models through a blood compartment (*9*). However, this model does not accurately describe the mass flow occurring in the human body, which starts with dietary intake followed by metabolism, transport, and elimination of the nutrients and its by-products. In the absence of such detailed representation, the generic human metabolic reconstructions have been used as a proxy for whole-body metabolism (*10, 11*). However, such approaches do not capture metabolic pathways that occur in parallel in multiple organs to give raise to known physiology, such as the Cori cycle.

To tackle these challenges, we developed a novel reconstruction paradigm yielding a molecular-level, organ-resolved, physiologically-accurate description of whole-body metabolism validated against current knowledge (Figure 1A). We demonstrate that WBM reconstructions can be converted into personalized WBM models through the integration of physiological, quantitative metabolomics, and microbiome data, thereby allowing the assessment of microbial metabolism on host metabolism in an organ-resolved, person-dependent manner.

**Figure 1:**
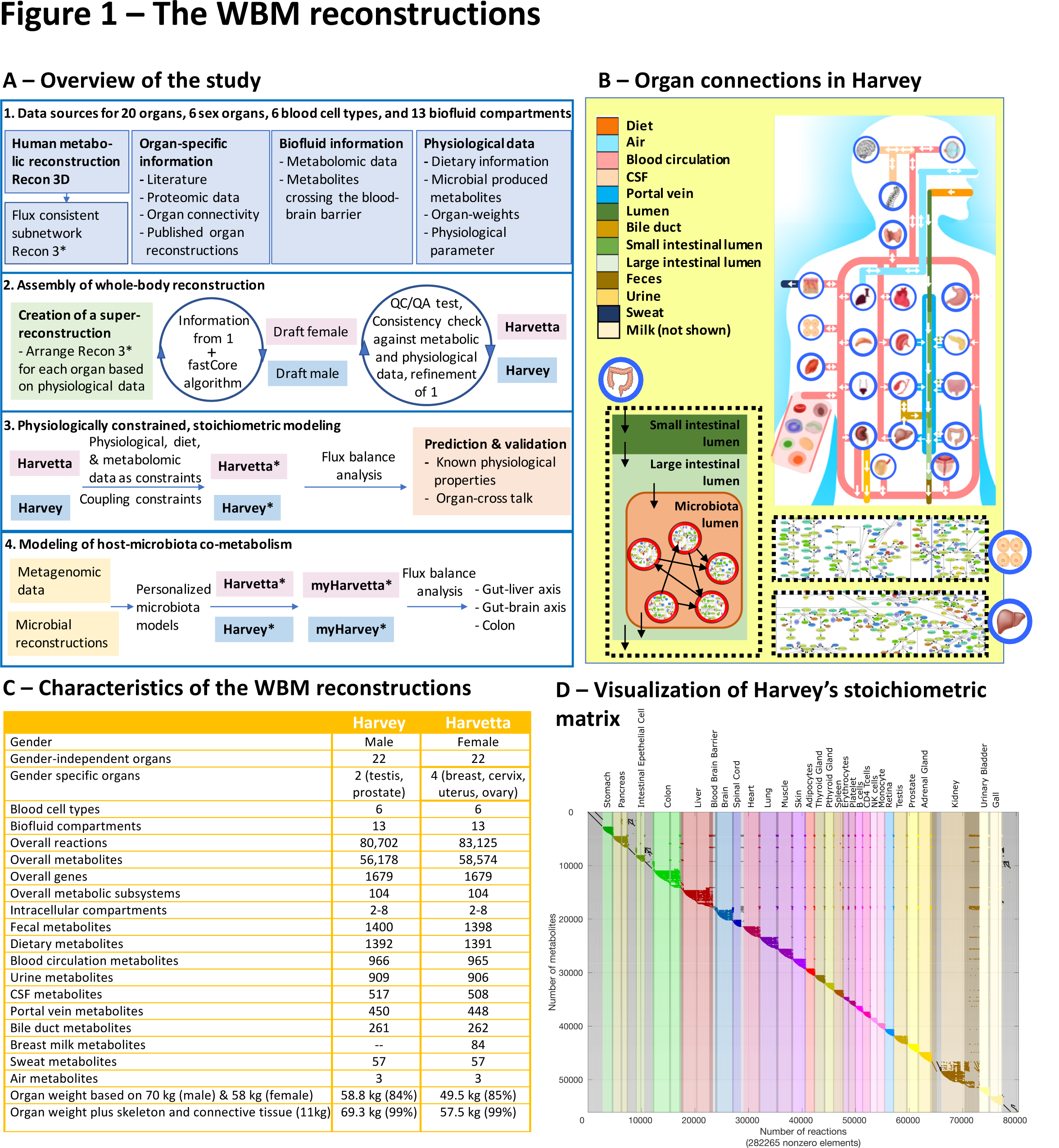
The whole-body metabolism (WBM) reconstructions. **A.** Overall approach to reconstruction and analysis of WBM model. Recon 3D has been assembled over the past ten years based on information from >2000 publications. The WBM reconstructions were assembled based on >600 publications and numerous omics data sets. myHarvey refers to the microbiome-associated WBM reconstructions. See Method section for more details. **B.** Schematic overview of the organs and their anatomical connections in Harvey. The arrows indicate the organ exchange with the biofluid compartments. **C.** Statistics of Harvey and Harvetta. **D.** Stoichiometric matrix of Harvey. Only non-zero entries are shown.

## Results

### Reconstruction of whole-body metabolism

We developed a novel, iterative approach assembling the organ-resolved WBM reconstructions based on an existing global reconstruction for human metabolism, Recon 3D (*3*), more than 600 literature articles and books, and omics data (Figure 1A). First, we generated a meta-reconstruction, comprised of 28 identical versions (32 for Harvetta) of Recon 3D, connected through respective biofluid compartments (Figure 1B) (Method section). Subsequently, each organ in the meta-reconstruction was tailored by removing reactions, which were found absent based on literature evidence or proteomic data (*12, 13*). The proteomic data provided evidence for the organ-specific protein expression and captured all organs except for the small intestine (Method section). The proteomic data and literature-derived information were used to define the core reaction set to be present in the different organs. The tailored meta-reconstruction and the core reactions set were subsequently subjected to a model extraction algorithm (*14*) to generate gender-specific draft WBM reconstructions (Figure 1A). Finally, as the algorithm determines one possible compact subnetwork for the meta-reconstruction and the core reaction set, we iteratively refined the content of the draft WBM reconstructions to ensure that the included reactions were consistent with the current knowledge about whole-body and organ-specific metabolism (Figure 1A).

Throughout this reconstruction process, we used established quality-control‐ and - assurance measures (Method section)(*15*). In total, the resulting WBM reconstructions account for more than 84% of the body weight, when excluding bones and connective tissue, which makes up another 15% of the body weight (*16*) (Method section). As one would expect, Harvetta contains more reactions than Harvey due to the four sex organs (Figure 1C, 1D). Both reconstructions shared 72,816 reactions, while 7,886 were unique to Harvey and 10,308 unique to Harvetta. When excluding the sex-specific organs, less than 4% of all network reactions were gender-specific, mostly involving alternative transport reactions (Method section). The resulting, gender-specific WBM reconstructions capture comprehensively human whole-body metabolism that is consistent with current knowledge.

### Organ atlas

To provide also organ-specific reconstructions, we extracted each organ from Harvey and Harvetta, while maintaining gender-specific reactions and the different blood compartments (Figure 1B, Figure 2) (Method section). We ensured that each organ followed the quality standard developed for metabolic reconstructions (*3, 15*), such as leak freeness, stoichiometric consistency, and realistic ATP yields from glucose under oxic and anoxic conditions (*3*). The organs contained on average 2,927±1,933 reactions, 2,063±1,081 metabolites, and 1,298±245 genes (Figure 2A). As one would expect, the organs with most reactions were liver, kidney, and colon. On average, over 70% of reactions in each organ were gene-associated, when excluding transport and exchange reactions (Figure 2A). The majority of the organ-specific reactions were in the liver, colon, small intestine, and the red blood cells (RBCs) About 10% of all metabolites were organ-specific metabolites (Figure 2B), which could be used as biomarker metabolites for organ-dysfunction. Again, most of these organ-specific metabolites were found in the colon (129), liver (74), and kidney (59). Notably, an additional 320 metabolites could be found only in two organs, potentially further expanding the set of organ-dysfunction markers. At the same time, each organ shared on average 29% of the metabolites in the blood compartment. The brain and the spinal cord shared 45 and 77%, respectively, of the metabolites with the CSF metabolic pool (Method section). Only 10% of the genes were core genes, which can be explained by the absence of the mitochondrial genes from the core set, as the RBCs are lacking this organelle. When excluding the RBC, an additional 142 genes were present in all remaining organs. There were 19 organ-specific genes in Harvey, with 11 and three being unique to the colon and small intestinal cells, respectively. For each gender and organ, we created a specific metabolic map, based on the map generated for Recon 3D, which can be used to visualize flux and omics data (*17*). Thus, the organ atlas represents a comprehensive set of manually curated, self-consistent organ-specific reconstructions that can be used for a range of biomedical applications (*18*).

**Figure 2:**
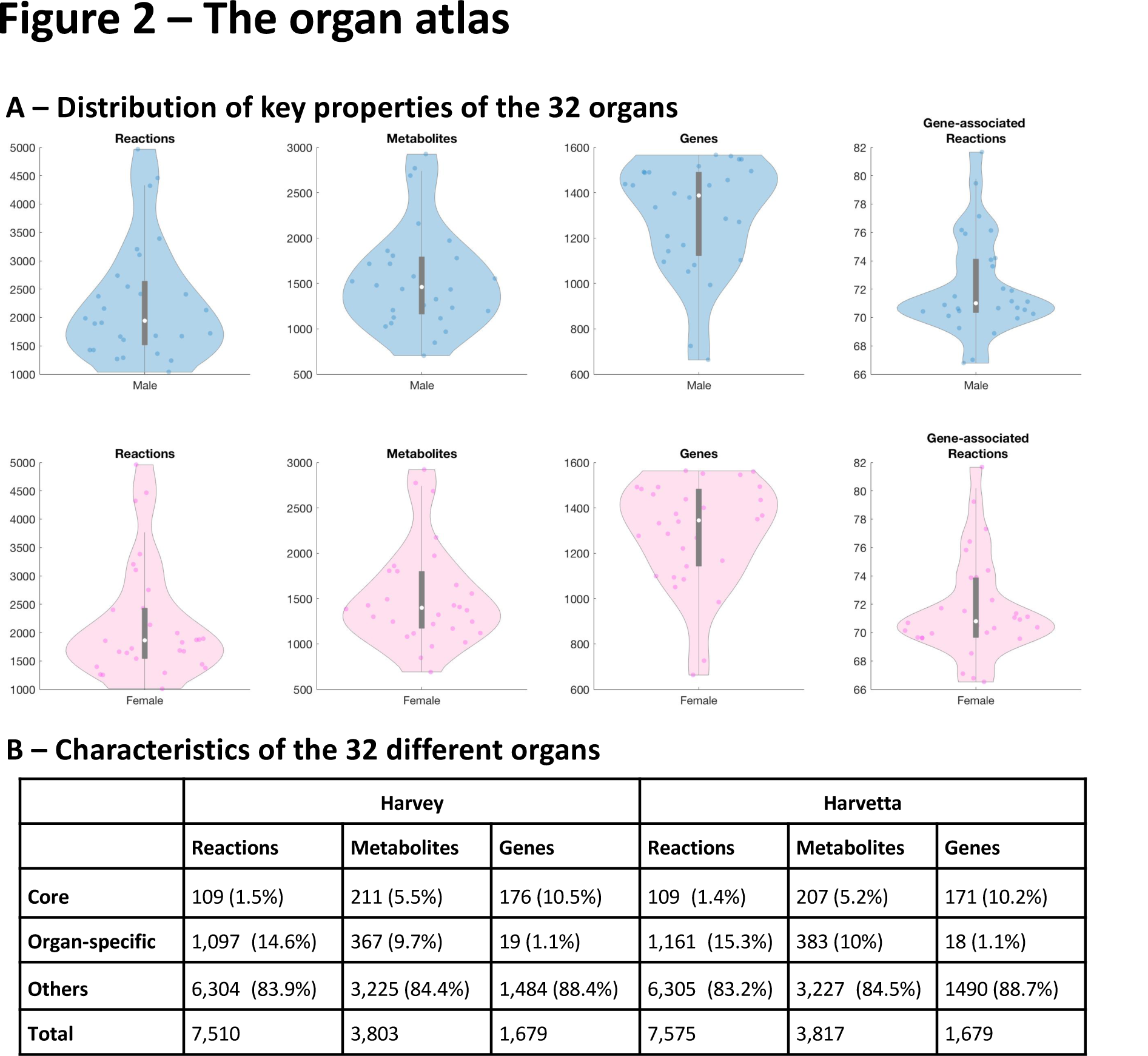
Characteristics of the organ compendium. **A.** Distribution of content male (blue) and female (pink) organ-specific reconstructions. **B.** Overall statistics of the organ compendium.

### Physiologically constrained, stoichiometric modeling

The conversion of a reconstruction into a model involves the definition of condition-specific constraints. To exploit the unique features of the WBM reconstruction, we constrained it using 15 physiological parameters (Figure 3A) (Method section), allowing us to quantitatively integrate metabolomics data (*19*) as modeling constraints. In particular, this allowed us to define how much of each metabolite an organ could take up, answering to the challenge to appropriately constrain organ-specific models. We also applied constraints for an average European diet (Figure 3B) (*20*). In total, 12.5% of the WBM model reactions had a constraint placed on its bounds, leading to a significantly reduced steady-state solution flux space. We will refer to this novel paradigm in constraint-based modeling as physiologically constrained, stoichiometric modeling (PCSM).

**Figure 3:**
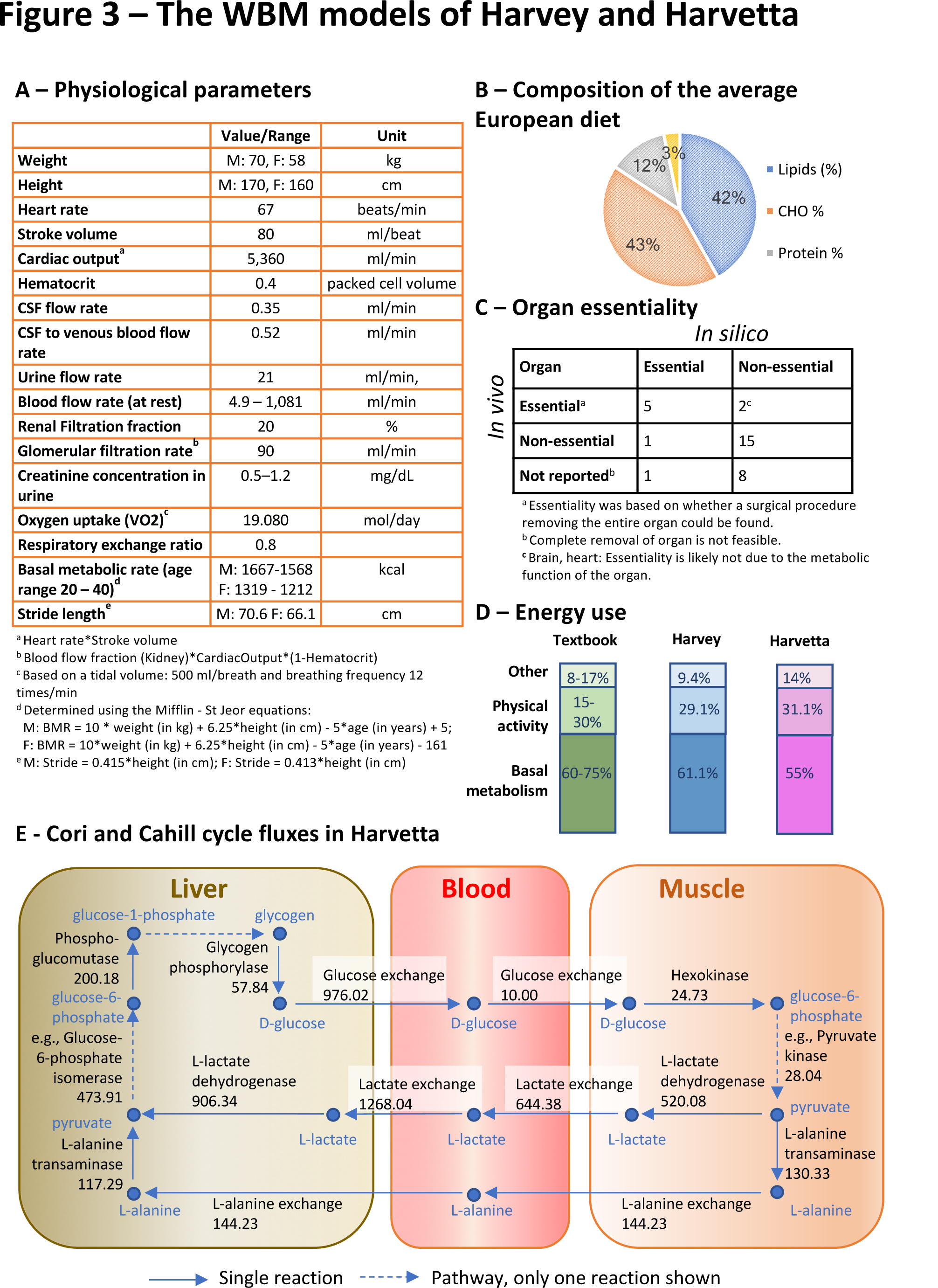
Integration of physiological parameters with the WBM reconstructions. **A.** List of physiological parameters used to constrain the reactions. The physiological values were retrieved for a reference man and woman (*16*). A complete list of constraints can be found in (Method section). **B.** Nutritional composition of the average European diet based on 2371.9 kcal. **C.** Predicted and in-vivo organ metabolic essentiality. **D.** Energy use is shown under the average EU diet. Textbook values are based on (*26*). **E.** Cori and Cahill cycle fluxes (in mmol/day/person). A single flux distribution, minimizing the Euclidian norm and subject to mass-balanced physiological constraints, for Harvetta is shown. Please refer to the Method section or <u>http://vmh.life</u> for reaction details.

### Basal metabolic flux

To represent the energy required to maintain the body's cellular function and integrity, we formulated a whole-body maintenance reaction, in which each organ biomass maintenance reaction is weighted based on their respective organ weight (Method section). This biomass maintenance assumes that the entire cellular proteome, transcriptome, and membranes are replaced once a day (*21, 22*). Consequently, we constrained the flux through this reaction be one and defined it basal metabolic flux (BMF), in analogy to the basal metabolic rate (BMR). We then calculated the overall energy (i.e., ATP) consumption of the WBM models, by minimizing the Euclidean norm (*8*). Assuming that the hydrolysis of one mole ATP yields 64 kJ (*23*), the overall energy consumption corresponded to 1,304 kcal and 1,460 kcal for Harvetta and Harvey, respectively. Harvey contains 11% more muscle, which has a higher ATP requirement in the muscle biomass maintenance reaction than the adipocytes. Our predicted energy consumption rate agreed well with the values obtained from the Mifflin-St Jeor equations, which estimate the BMR (Figure 3A) based on gender, weight, height, and age, and explain about 70% of the observed variability in individuals (*24*). Hence, the BMF in the WBM models recapitulates the basal metabolic function of the whole body. Physical activity consumes the remaining energy and nutrient resources provided by the diet.

To further validate the predictive accuracy of Harvey and Harvetta, we computed each organ's essentiality for the BMF under the given physiological and dietary constraints. The predictions were consistent with which organs can be fully surgically removed (Figure 3C). We then minimized the Euclidean distance in the simulations to determine the resting brain energy consumption in Harvetta (10.21 mole/day/person) and Harvey (11,31 mole ATP/day/person). These predicted values are about half of the reported brain consumption 120 g of glucose per day and brain (*25*), which corresponds to 20.65 mole ATP/ person/day when assuming conversion of 1 mole of glucose into **31** moles of ATP. Hence, further parameterization of the energy consumption for the non-resting brain will be necessary. Overall, these examples demonstrate that the WBM models capture emergent, complex, and gender-specific phenotypes.

### Energy homeostasis

We investigated the capability of Harvey and Harvetta to either store fat in the fat cells, or adipocytes, or to use the dietary energy for muscle work. Note that we did not change any physiological parameters; hence, this associated physical activity corresponds to slow walking. The maximal possible fat storage flux of Harvey was 6% higher than that of Harvetta (81.76 mmole triglyceride/day/person vs. 77.15 mmole triglyceride /day/person), while the maximal possible muscle energy flux was in 7% higher in Harvetta (45.11 mole ATP/day/person vs. 48.35 mole ATP/day/person). The reason for this counter-intuitive result is that Harvey has 11% more muscle tissue (40% vs. 29%), but 11.3% fewer adipocytes (21.4% vs 32.7%), thus leading to an increased muscle maintenance cost and reduced comparative ATP hydrolysis capability.

The daily energy requirement consists of the BMR and activity-induced energy expenditure (Figure 3) (*26*). The predicted maximal possible muscle energy flux values correspond to 690 kcal for Harvey and 740 kcal for Harvetta. Consistent with the current understanding of energy use (*26*), the predicted muscle energy consumption flux corresponded to about 30% of the dietary energy intake, while the BMF represented about 55-61% of the dietary intake (Figure 3). The remaining 9-14% of the dietary caloric intake were unused by in the computed flux solution, partially due to the lower brain energy consumption in the WBM models. *In-vivo*, 8-17% of the dietary energy is used for food-induced thermogenesis and other non-metabolic processes (*26*).

To put the predicted activity-induced energy expenditure into context, we determined the corresponding number of steps, assuming a gross energy cost of 3 J/kg/m (*27*) and a particular stride length (Figure 3A) (Method section). Harvey would have to walk 9,701 steps and Harvetta 11,752 steps to utilize the predicted muscle energy. These numbers are in agreement with the popular recommendation to walk a 10,000 steps a day (*28*). To reduce their weight, Harvey and Harvetta would have to consume more energy, which could be achieved *in-silico* by increasing the heart rate and oxygen consumption, thus providing more nutrients and oxygen to the organs. In an agreement, daily physical activities corresponding to about 11,000–13,000 steps have been recommended to prevent weight gain (*28*). Harvetta's higher maximal possible flux through the muscle ATP demand is consistent with suggested gender-specific differences of the physical activity threshold (*29*), and knowledge that women store more fat than men (*30*).

Overall, this analysis demonstrates that the WBM models capture known physiological, gender-specific characteristics, which also arise from different organ distributions, particularly the adipocyte-muscle ratio.

### Host-microbiota metabolic interactions

The microbiota, directly and indirectly, influences host metabolism (*31*), including the brain (‘gut-brain axis’), the liver (‘gut-liver-axis’), and the colon. To assess the effect of individual microbiota on host metabolism, we mapped strain-resolved metagenomic data for 149 healthy individuals provided by the Human Microbiome Project (HMP) Consortium (*32*) onto a resource of gut microbial metabolic reconstructions, AGORA (*5*), as described previously (*33*). The microbiome models captured 917% of the relative phylum-level abundance. We then joined each microbiome reconstruction with either Harvey or Harvetta (Figure 1B) and personalized them using the weight, height, and heart rate of each individual, resulting in personalized microbiome-associated WBM models (Figure 4A) (Method section). Personalized germ-free WBM models were obtained by setting microbiota community biomass reaction flux to zero.

**Figure 4:**
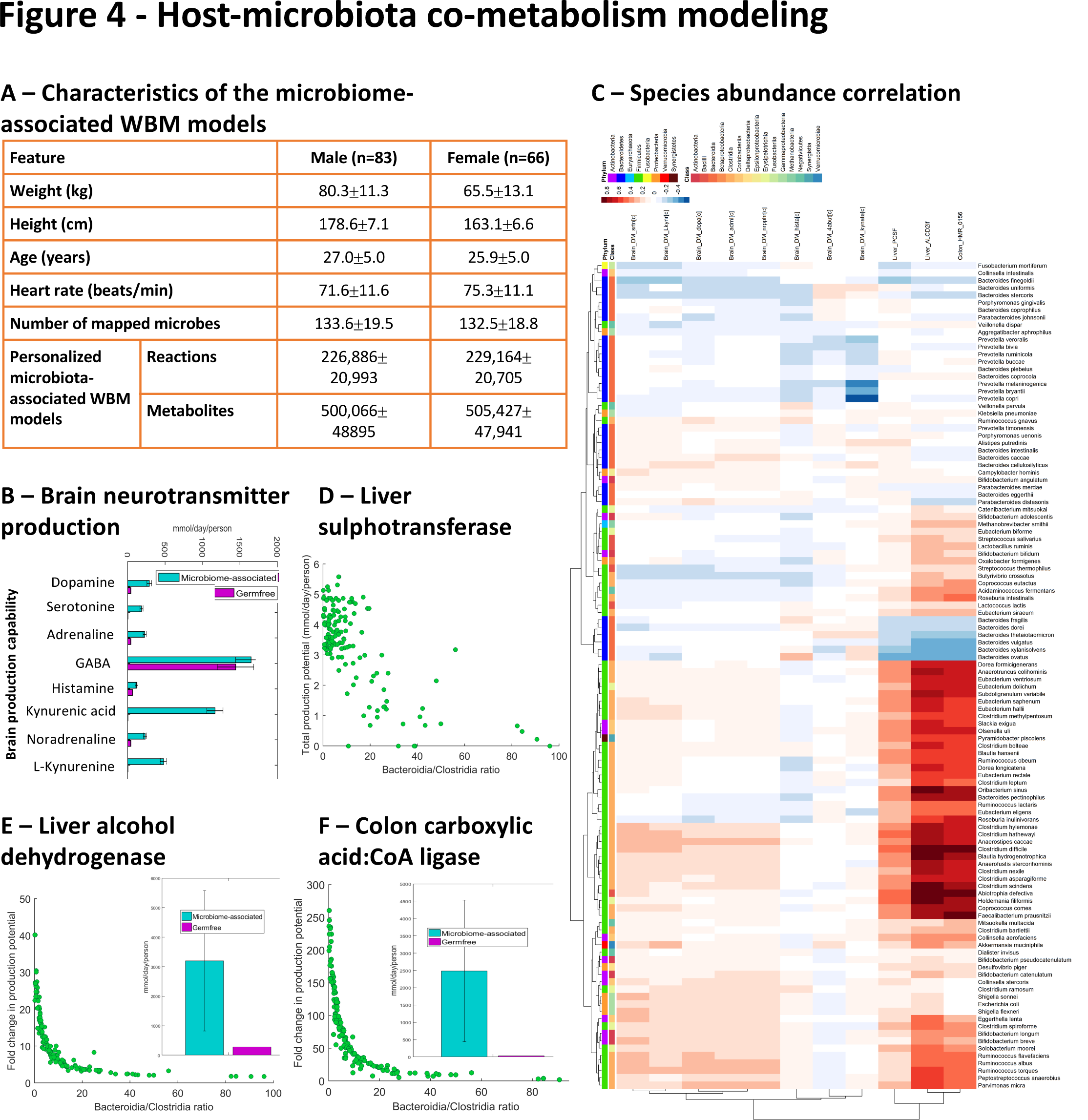
Host-microbiota co-metabolism in 149 personalized WBM models. **A.** Characteristics of the microbiome-associated WBM models. **B.** Average brain biosynthesis potential for eight neurotransmitters in the personalized WBM models with and without microbiome (germfree). **C.** Correlations between species-level abundances and fold changes (microbiome-associated vs. germfree) for flux through 11 objective functions. **D-F.** Bacteroidia/Clostridia ratio against maximal flux (mmol/day/person) against (relative abundance) through the different reactions. Inlet: Average maximal reaction flux in microbiome-associated and germfree WBM models.

The gut microbiota produces various neurotransmitters and the metabolic precursors (*31*). To determine the microbiota contribution to the brain neurotransmitter availability, we first maximized for the accumulation of eight prominent neurotransmitters in the germ-free personalized WBM models and observed substantial inter-individual variation. The addition of the individual microbiomes led to a substantial increase in neurotransmitter availability with a high inter-individual variation (Figure 4B, 4C) that could neither be explained by the presence of a single species or genus (Figure 4B) nor by the provided meta-data. Consistently, in the microbiome models without the host, a variety of strains across multiple taxa were found to produce the amino acid precursors, as well as the neurotransmitters GABA and histamine (Supplemental Figure S2). Thus, WBM modeling captures the known role of the gut microbiota as an additional organ influencing brain metabolism (*31*).

The human gut microbiota can also influence host drug metabolism (*34*). One prominent example is the production of p-cresol, e.g., by *Clostridium difficile* (*Clostridioides* genus), which competes with pharmaceuticals, such as acetaminophen, for sulphonation in the liver and thus interferes with drug detoxification (*34*). The resulting product, p-cresol sulfate, can lead to kidney impairment (*35*). In microbiota with low *Clostridioides* genus abundance, the genus directly correlated with the maximally possible flux through the liver sulphonation reaction, while at higher abundances, substrate availability was limiting for p-cresol production and subsequent sulphonation (Supplemental Figure S3). Liver sulphonation also inversely correlated with the abundance of the Bacteroidia class (Figure 4D). Thus, WBM modeling allows for the prediction of individual-specific p-cresol sulfate levels. p-cresol sulfate has been proposed as a predictive biomarker for drug detoxification (34 and mortality in chronic kidney disease (*35*).

Ethanol directly causes liver toxicity and compromises intestinal barrier function (*36*). Its product acetaldehyde is cytotoxic and carcinogenic (*36*). The flux through liver alcohol dehydrogenase was increased by a fold change of 11.48.5 compared with germfree (Figure 4E). This fold change in flux strongly correlated with species belonging to the Clostridia class and negatively correlated with representatives of the Bacteroidia class (Figure 4E). Species that showed the highest correlations with alcohol dehydrogenase flux included *Clostridium scindens*, *Blautia hydrogenotrophica*, and the well-known pathobiont *Clostridium difficile* (Figure 4C). This example further illustrates that WBM modeling may have valuable applications for *in silico* clinical trials with its ability to predict an individual-specific toxicity at organ-level.

Butyrate, which is mainly produced by gut microbes in the Clostridia class, serves as the main energy source for the colonocyte (*37*). We computed the flux through the butyrate-CoA ligase in the colonocyte. As expected, the butyrate-CoA ligase flux was significantly increased in the presence of the microbiome by a fold change of on average 78.764.7 (Figure 4F). The butyrate-CoA ligase flux strongly correlated with the abundance of Clostridia (Figure 4F) and with known butyrate producers, such as *Faecalibacterium prausnitzii*, *Butyrivibrio crossotus*, and *Subdoligranulum variabile* (*37*) (Figure 4C). In conclusion, Harvey captures the individual-specific cross-feeding of butyrate to the human host.

Taken together, using the personalized microbiome-associated WBM models, we demonstrated that the gut microbiome increases the neurotransmitter production potential in the brain, modulates detoxification enzymes in the liver, and provides butyrate to the colonocyte with high inter-individual variability.

## Discussion

We presented a gender-specific, organ-resolved, molecular-level, physiologically accurate reconstruction of human whole-body metabolism. The underlying detailed description of human metabolic pathways has been developed over the past decade based on more than 2000 literature articles and books (*3*), and provided an indispensable foundation for Harvey and Harvetta. Organ-specific metabolism (Figure 2) was based on and validated against >600 published studies and books, and accounts for comprehensive proteomic and metabolomics data (Method section). Known inter-organ metabolic interaction, as illustrated with the classical examples of the Cori and Cahill cycle, as well as organ-essentiality (Figure 3) were captured with the WBM reconstructions, as were whole-body functions, such as BMR and the energy use (Figure 4). Using the microbiota-associated WBM models, we could demonstrate the microbial influence on different organs (Figure 4) consistent with knowledge mainly derived from animal studies. Finally, personalized versions of the WBM models reflect inter-individual variability in metabolic response to varied physiological parameters and external clues consistent with phenomenological observations. Taken together, the WBM reconstructions represent a molecular-level description of organ-specific processes built on current knowledge and underpinned on basic physical principles.

The creation of organ-specific metabolic reconstructions is challenging despite the myriad of omics data and sophisticated algorithms (*4*). We tackled this challenge by an iterative approach combining extensive literature review, omics data, and high-performance computing (Figure 1). Moreover, the inclusion of biofluids in the WBM enabled the integration of metabolomics data, while the use of microbiome information and dietary information increased the comprehensiveness of the WBM. Hence, the metabolic complexity captured in Harvey and Harvetta could not have been achieved using an organ-by-organ level reconstruction approach. This novel reconstruction paradigm could present a blueprint for other multi-cellular model organisms, such as the virtual physiological rat (*38*).

Harvey and Harvetta capture anatomically-accurate connections between organs. Current approaches have either assumed the whole-body human metabolic network without organ-boundaries (*10, 18*) or simplified it (*9*), which ultimately limits its general usability and predictability. Here, in contrast, we consider individual-level physiological, nutritional and microbial parameters for personalization of the WBM models and provide a novel tool to study the inter-person variability in physiological processes as well as drug metabolism. Personalization of computational human models is a requirement for enabling *in-silico* clinical trials (*39*). Importantly, Harvey and Harvetta can be expanded to include other cellular (e.g., signal transduction and regulation) as well as whole-body level processes (e.g., drug metabolism) by serving as a docking station. Using metabolic models as a platform for integration of other types of models have been already demonstrated, most notably the whole-cell model of *Mycoplasma genitalium* (*40*), and could now be expanded to whole-body human metabolism.

The PCSM approach allows the integration of physiological parameters and quantitative metabolomics data to constrain the organ-uptake rates to physiologically relevant values and thereby limit the achievable intra-organ metabolic flux states. Our approach has a reasonable computing time as only one linear (or quadratic) programming problem has to be solved. In contrast, hybrid modeling approaches, such as the ones integrating physiology-based pharmacokinetic modeling with genome-scale models (*41*), require solving the considered metabolic models (e.g., hepatocyte model) at each time step. Consequently, solving hybrid models is computationally expensive. The processes in the human body are intrinsically dynamic, but not all biomedical questions require the dynamical consideration. In those cases, Harvey and Harvetta are valuable, time-efficient alternatives, and they capture human metabolism at a more comprehensive level than currently practical for dynamical (hybrid) models.

In addition to our genome, the diet and the microbiome contribute to inter-individual variations in disease development and progression (*42*). Computational modeling accounting for these environmental factors has been demonstrated (*11, 43*) but is hampered by the lack of a molecular-level, organ-resolved description of human metabolism. For instance, the gut microbiota has been suggested to modulate human metabolism on an organ-level but the underlying pathways are typically elucidated using animal models (*44*). Harvey and Harvetta enable such analysis *in-silico* and a comparison with their germ-free counter-part. This capability provides an unprecedented opportunity to develop novel, mechanism-based hypotheses how the gut microbes, individually and collectively, modulate human metabolism. While these hypotheses will require experimental validation, the WBM models permit testing of experimentally-derived hypotheses and prioritize subsequent experimental studies, thus accelerating knowledge creation through a combined *in-silico – in-vitro* approach.

Taken together, Harvey and Harvetta represent a significant step towards the “virtual human” envisioned in the Tokyo declaration (*6*).

## Methods

### Reconstruction details

In this part, we describe the reconstruction approach and the information used to generate the whole-body metabolic (WBM) reconstructions, Harvey and Harvetta. The simulation-relevant information and methods can be found in the section “Materials and Methods – Simulation details”.

#### Whole-body metabolic reconstruction approach

As a starting point for the WBM reconstructions, we used the global human metabolic reconstruction, Recon 3D (*3*), which accounts comprehensively for transport and biochemical transformation reactions, known to occur in at least one cell type. Recon 3D can be obtained Recon 3D from the Virtual Metabolic Human database <u>(http://vmh.uni.lu/#downloadview)</u>. While the reconstruction of Recon 3D consists of 13,543 reactions, 4,140 unique metabolites, and 3,288 genes, the flux and stoichiometrically consistent global metabolic model (Recon 3D model) contains 10,600 reactions, 5,835 non-unique metabolites (2,797 unique metabolites), and 1,882 unique genes. We removed from Recon 3D model those reactions, and metabolites, involved in protein and drug. We then removed flux inconsistent reactions (i.e., those reactions that did not admit non-zero net flux). The resulting metabolic model, Recon 3*, contained 8418 reactions, 4489 (non-unique) metabolites, 2053 transcripts, and 1709 genes. In Recon 3*, each enzyme-catalyzed reaction or transport reaction is associated with the corresponding gene(s) that encode the protein(s). These so-called gene-protein-reaction associations (GPRs) represent through Boolean rules (‘AND, ‘OR’) isozymes or protein complexes.

### Setup of the whole-body, gender-specific meta-reconstructions

The central idea was to generate a human organ-resolved whole-body model that was built not by connecting the separate metabolic subunits, i.e., the individual organs, but which would emerge as one functional, self-consistent whole-body metabolic system from the metabolic capabilities and the known interactions between the organs.

Therefore, we considered 20 organs, six sex-specific organs, and six blood cells were considered (Table 2). For simplicity, we refer to all of these as organs in the following. We defined 13 biofluid compartments to be considered in the WBM reconstruction (Figure 1B, main text, Table 1). For each organ, transport reactions from the extracellular compartment ([e]) to the blood compartment ([bc]) were added to Recon 3*. Additional transport reactions were added to those organs that are connected to a third, or forth, biofluid (e.g., liver, Figure S1). For organs, which can only take up from or secrete into a particular biofluid (see arrows in Figure 1B, main text), the reaction directionality was set accordingly. The transport mechanism was always through facilitated transport, which assumes that the metabolites can be transported from the biofluid to the interstitial fluid surrounding the organ cells is driven either by concentration difference (diffusion) or pressure difference (bulk flow). Each reaction in Recon 3* and the newly added transport reactions received a suffix corresponding to one organ (Table 2). Numerous organs are known to store metabolites, e.g., liver. We included sink reactions for stored metabolites in the corresponding organs (Table S3). We then joined all organ-named Recon 3* versions to create two meta-reconstructions, which represent the organ connectivity in an anatomically accurate manner for the female and male metabolism (Figure 1B, main text). We added diet uptake reactions for all metabolites with defined exchange reactions in Recon 3*, as well as transport reactions along the gastrointestinal tract (Figure 1B, main text, Table 1) to the meta-reconstructions. Both, small intestinal epithelial cells (sIEC) and colonocytes were allowed to take up metabolites from their corresponding luminal compartment (Table 1) and could secrete some metabolites into the lumen (Table 3). Gut microbes are known to produce in the gastrointestinal tract valuable metabolic precursors to the human host. To enable the uptake of such metabolites by the small intestinal cells and the colonocytes, we added sink reactions into the luminal compartments of the meta-reconstructions.

**Table 1:** Biofluid compartments and the connected organs.

| Biofluid compartment | Abbreviation | Connected organs |
| --- | --- | --- |
| Diet | [d] | - |
| Lumen | [lu] | - |
| Lumen, small intestine | [luSI] | Small intestinal cells |
| Lumen, large intestine | [luLI] | Colonocytes |
| Feces | [fe] | - |
| Blood, circulation | [bc] | All but brain, spinal cord |
| Blood, portal vein | [bp] | Liver, colonocytes, small intestinal cells, pancreas, spleen |
| Bile duct | [bd] | Liver, gallbladder |
| Cerebrospinal fluid | [csf] | Brain, spinal cord |
| Urine | [u] | Kidney |
| Sweat | [sw] | Skin |
| Breast milk (female only) | [mi] | Breast |
| Air | [a] | Lung |

**Table 2:** List of organs in Harvey and Harvetta.

| Organ name | Organ prefix |
| --- | --- |
| Adipose tissue | Adipocytes_ |
| Adrenal gland | Agland_ |
| B-cells | Bcells_ |
| Brain | Brain_ |
| Breast | Breast_ |
| CD4+ T-cells | CD4Tcells_ |
| Cervix | Cervix_ |
| Colon | Colon_ |
| Gallbladder | Gall_ |
| Heart | Heart_ |
| Kidney | Kidney_ |
| Liver | Liver_ |
| Lung | Lung_ |
| Monocytes | Monocyte_ |
| Muscle | Muscle_ |
| Natural killer cells | Nkcells_ |
| Ovary | Ovary_ |
| Pancreas | Pancreas_ |
| Platelet | Platelet_ |
| Prostate | Prostate_ |
| Pthyroid gland | Pthyroidgland_ |
| Red blood cell | RBC_ |
| Retina | Retina_ |
| Spinal cord | Scord_ |
| Small intestine | sIEC_ |
| Skin | Skin_ |
| Spleen | Spleen_ |
| Stomach | Stomach_ |
| Testis | Testis_ |
| Thyroid gland | Thyroidgland_ |
| Urinary bladder | Urinarybladder_ |
| Uterus | Uterus_ |

**Table 3:**
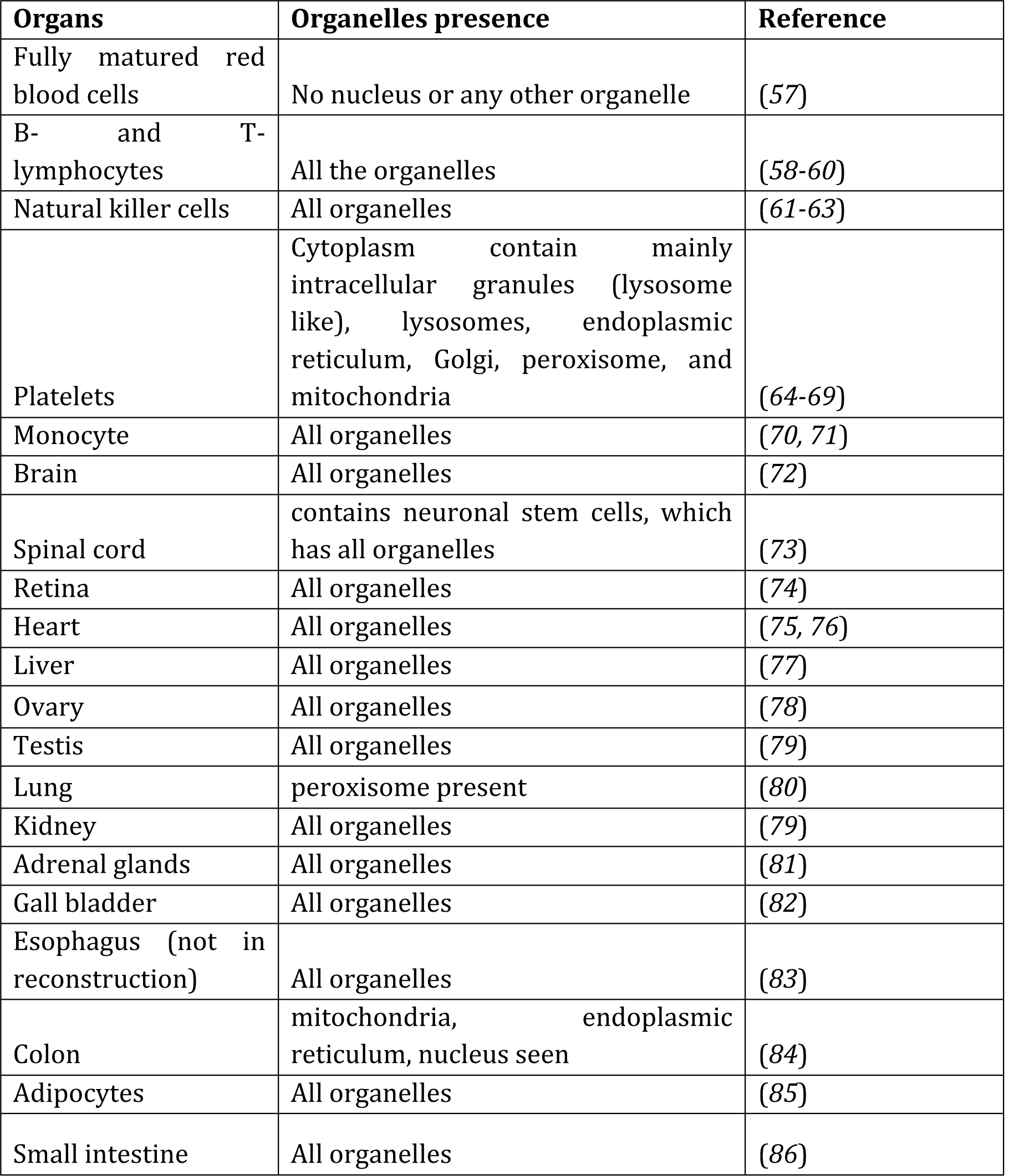
List of evidence for organelles in the different organs.

## Anatomically accurate organ connectivity

The dietary input compartment represents the exchange medium consisting of all the dietary ingredients that the human body can consume. The dietary inputs from the [d] enter the gastrointestinal lumen represented by [lu] in Harvey. The lumen compartment set up in the model represent the gastrointestinal lumen which is unidirectional and exits into the fecal excretion compartment [fe]. The fecal excretion compartment represents the excretory end-products comprising the undigested and unabsorbed part of the dietary input. In Harvey, the gastrointestinal lumen compartment is further divided into the small intestinal lumen and the large intestinal lumen. While the gastrointestinal lumen receives the diet input, the small intestinal lumen receives metabolite drainage from the gallbladder (via the bile duct), pancreas, and the small intestine. The large intestinal lumen is specific for the large intestine receiving metabolites only from the colon. The portal venous blood receives metabolites from the colon, small intestine, spleen, pancreas, and gallbladder, which finally drains into the liver for their further metabolism, and for exposure to the systemic circulation. The bile duct is a special compartment, which is specific for liver and gallbladder. Bile is synthesized in the liver and stored in the gallbladder (*45*). From the liver, bile flows into the bile duct. The flow of bile into the small intestine via the bile duct depends on the Sphincter of Oddi that closes during the inter-digestive period, increasing the pressure in the bile duct, and resulting in backflow of bile into the gallbladder, where it is further concentrated. During the digestive phase, the Sphincter of Oddi opens causing the concentrated bile flow into the small intestinal lumen to aid in digestion (*46*).

The systemic blood circulation is represented by the circulatory blood in Harvey, which provides oxygen and nutrients to all organs. Since the brain and the spinal cord are specific in its metabolite exchange, we introduced the blood-brain barrier and cerebrospinal fluid as extracellular compartments. The blood-brain barrier selectively allows the exchange of metabolites, to and from the brain (explained above), and the cerebrospinal fluid receives metabolites from the brain, finally draining into the circulatory blood compartment (Figure 1, main text). The lung takes up oxygen from the environment and gives out carbon dioxide, which is captured as [a] in the WBM reconstructions. Finally, the urine compartment contains filtered metabolites from the kidney, for final excretion. These extracellular compartments signify precise anatomical barriers of the organs or cells of the human body.

### Generation of draft gender-specific WBM reconstructions from the meta-reconstructions

The first step was to arrange 28 times (32 for Harvetta) Recon 3* in an anatomically correct manner as described above (Figure 1 B, main text). The resulting meta-reconstruction was then to tailored to represent organ-specific metabolism. To achieve this, we used the fastCore algorithm (*14*). Briefly, fastCore takes as input a metabolic reconstruction and a set of core reactions, known to be active in the network, to identify a flux-consistent subnetwork containing all core reactions (as long as they are flux consistent) and a minimal number of additional reactions. To define the core reactions for each organ, we used two comprehensive proteomic data sets (*12, 13*), providing information on the organ-specific protein expression of all 1691 but 18 genes included in Recon 3*. No protein expression data was available for the small intestinal cells. We required that at least one reaction associated with the organ-specific expressed gene was present in the corresponding organ. Reactions from four published organ-specific reconstructions (i.e., red blood cell (*47*), adipocyte (*9*), small intestine (*48*), and liver (*49*)), were matched to the name-space of Recon 3* and also added to the core reaction set (see also below). Additionally, we also incorporated information from more than 500 literature resources for the presence or absence of metabolic and transport reactions, genes, or pathways in all organs (see below). Particular emphasis was placed on the metabolism occurring in skeletal muscle, skin, spleen, kidney, lungs, retina, heart, and brain (see below for details on these organs). In the case of absence, the corresponding organ-specific reactions were set to have a lower and upper bound of 0, thus effectively removing these reactions from the organ. Note that this input data was not gender-specific, due to the absence of corresponding data on an organ-specific larger scale. We considered for each gender two different scenarios and set up constraints on the meta-reconstructions accordingly. First a feeding condition was set up, in which all dietary exchange reactions were open (i.e., the lower bound was set to ‐inf and the upper bound was set to zero) and the storage of metabolites in the various organs was enabled (lower bound on the sink reactions were set to be zero and the upper bound was set to be +inf). Second, a fasting condition was set up, in which all dietary uptake reactions were closed (lower and upper bound were set to zero) but access to stored metabolites from the different organs was enabled (sink reactions had a lower bound of ‐inf and an upper bound of zero). To ensure that metabolites found in different biofluids based on metabolomics data (see below) would also be present in the WBM reconstructions, we added demand reactions for all those metabolites to the biofluid compartments of the four meta-reconstructions and added them to the core reaction set. By doing so, we enforced that at least one organ could produce a given metabolite, or that it would be taken up from the diet (and the lumen). Using these four differently setup meta-reconstructions along with the input information (the corresponding core reactions for each setup) and the fastCore algorithm, we generated four flux-consistent subnetworks. We then removed the demand reactions for the biofluid metabolites, defined all reactions present in each subnetwork to be the core reaction set for the given setup, and repeated the extraction of flux consistent subsets from the meta-reconstructions. By doing so, we also enforced that at least one reaction could either transport or excrete a given biofluid metabolite or that it is catabolized in at least one organ.

Finally, we joined for each gender the fasting and the feeding subnetworks. The rationale for having the feeding and the fasting condition is that the human body can fast overnight, and thus the WBM reconstructions should capture this capability regarding catabolic as well as anabolic metabolic reactions. Note that the WBM reconstructions are not able to starve, as this would require the degradation of muscle proteins, which we did not explicitly capture in Recon 3*, and thus in the WBM reconstructions.

fastCore adds a minimal number of additional reactions to the core reactions to form the flux consistent, compact subnetwork. Hence, the added reactions represent hypotheses of which reactions and pathways would be needed to make the subnetworks flux consistent, given a set of core reactions. It does not mean that the proposed solution is biologically relevant. Consequently, after the generation of the male and female draft WBM reconstructions, we manually inspected that the added reactions were consistent with the current knowledge about organ-specific metabolism. The core reactions and the absence of organ-specific reactions were updated based on literature evidence, and the subnetwork generation was repeated. Overall, due to the complexity of the WBM reconstructions and the large number of reactions (more than 80k, Figure 1B, main text), we iterated this process more than 100 times, each time focusing on different organs or pathways (spanning multiple organs).

Moreover, WBM model predictions were compared with organ-specific data and known whole body metabolic functions (e.g., multi-organ metabolism of glucose (Cori cycle), amino acid cycle (Cahill cycle)). In this step, we added missing reactions to the gender-specific draft reconstructions. During the entire reconstruction process, we performed the same quality control and assurance tests as defined for metabolic reconstructions (*15*). At its end, the resulting WBM reconstructions represent human physiology and organ-specific metabolism to the best of our knowledge.

#### Details on the core reaction sets

For each organ, a core reaction set was defined that included organ-specific (i) protein information from the human proteome map (HPM) (*12*) and the human protein atlas (HPA) (*13*), (ii) extensive manual curation of organ-specific literature, and (iii) reactions presence in four published tissue-specific organs, i.e., red blood cell (*47*), adipocyte (*9*), small intestine (*48*), and liver (*49*).
i. **Proteomic data** The map of the human proteome (HPM) (*12*) provides protein information for 17,294 genes that accounted for 84% of the protein-encoded part of the human genome. These proteins were obtained from normal histological tissue samples, accounting for 17 adult tissue types, from three deceased individuals. We queried the database (10/12/2015) for all genes present in Recon 3*. For 1678/1709 Recon 3* genes/ proteins (98% coverage) to obtain their distributions in the 23 tissue types. The obtained protein expression data were scaled to range from 0 to 1. Only those proteins with an expression level of greater or equal to 0.2 were assumed to be expressed in an organ. Moreover, to complement the HPM resource, we used HPA (*13*). The protein expression data for the normal tissues was downloaded and sorted according to the tissue type. The proteins in the human protein atlas have been analyzed with a single antibody, and the expression levels have been reported based on the antibody staining (*50*). The expression levels depend on the quality of the antibody used (*50*). Hence, we considered only those proteins, which had a high and medium expression level for a given tissue/organ. For the brain, we combined the protein information of cerebellum, cerebral cortex, hippocampus, and lateral ventricle. The Ensembl gene IDs were converted to their corresponding EntrezGene IDs using the BioMart tool from Ensembl (*51*). We used the GPR associations given in Recon 3* to identify reactions to be present in the core set. Note that in the case of protein complexes, the presence of one of the proteins from the complex was deemed sufficient to require the presence of at least one reaction in the WBM reconstruction. After obtaining the reaction list we incorporated it into the core set for each respective organ reconstruction.
ii. **Literature-based curation of WBM reconstruction content** We performed manual curation of reactions, genes, and pathways for all included organs but focused in particular depth on the metabolism occurring in eight organs, which contribute substantially to inter-organ metabolism (*52*). These organs were: skeletal muscle, skin, spleen, kidney, lungs, retina, heart, and brain. We followed the bottom-up, manual reconstruction approach established for metabolic reconstructions (*15*). We collected information on the pathways/reactions that are absent across organs. Primary literature articles, review articles, and books on organ-specific metabolism were thoroughly studied to derive the pathway information.

## Definition of metabolic units

In Recon 3*, reactions are grouped into distinct sub-systems, which represent the overall metabolic process, e.g., glycolysis/gluconeogenesis. However, such broad categorization is of limited use, when building a cell or tissue specific metabolic network. A particular metabolic pathway may be selectively active across multiple tissue types. A typical example is the degradation of branched chain amino acids, i.e., valine, leucine, and isoleucine. While the skeletal muscle possesses the entire pathway (*53*), only the first two reactions of this pathway have been reported to be occurring in the kidney (*54*). Additionally, in case of fatty acid oxidation reactions, certain tissues can selectively oxidize fatty acids, e.g., kidney actively oxidizes octanoic acid, at similar rates as palmitic acid (*55, 56*). Therefore, it is essential to formulate 'metabolic units' that not only account for the Recon 3* sub-systems but are also metabolite specific.

We categorized individual metabolic and transport reactions in Recon 3* as metabolic units. Each metabolic unit contains three components, (i) major metabolic pathway, (ii) product formed, and (iii) cellular location, which represents the first and the last reaction steps of a particular pathway, respectively.

The reaction content of Recon 3* was classified into 427 metabolic units when only the major metabolic pathway was considered and the cellular compartments ignored. When the whole metabolic unit along with all its components was taken into account, 5637 metabolic units resulted.

Usage of the metabolic units greatly accelerated the reconstruction process as they account for individual metabolite-specific pathways as well as key enzymes of the biochemical pathways, as they are frequently reported and referred to in the biochemical literature. This literature information was translated into occurrence and non-occurrence of metabolic units. Additionally, we noted tasks that an organ can carry out (e.g., storage of glycogen) or the inability to carry out a particular task (e.g., storage of vitamins occurs only in a limited number of organs), leading to the formulation of organ-specific metabolic objectives.

## Presence of cellular organelles in organs

Within the blood tissues, differences in the presence of cellular organelles exist between fully matured red blood cells and others. For instance, fully matured red blood cells are devoid of a nucleus or any other organelles (*57*). The other hematopoietic cells, such as the B-lymphocytes, T-lymphocytes, natural killer cells, and monocytes contain all the cellular organelles (*58, 61, 70*). Therefore, we performed a thorough manual search and obtained a definitive occurrence of cellular organelles for 19/32 organs from the literature (Table 3). This search was important to represent the organ's metabolic capabilities accurately in the WBM reconstructions. For the remaining organs, no information could be found in the literature.

## Nutrient storage within organs

Maintenance of certain metabolite pools and metabolite storage as a reserve for energy demands within the cells has been reported to be crucial for maintaining the organ-specific functions. Typically, these are the glycogen storage in liver and skeletal muscle (*45*), or fatty acid storage in the adipocytes (*87*). During periods of fasting, liver glycogen serves to maintain the blood glucose levels. Additionally, triglyceride stores in the adipocytes are broken down to supply fatty acids to skeletal muscle and heart to serve as an energy resource (*88*). A thorough manual search of the storage capacity for dietary nutrients by various organs was performed. Known storage capacities were represented by adding specific demand/sink reactions (Supplementary Table S5) to the corresponding organs, and the core reaction sets. The demand reactions serve as cellular demand or usage of the metabolites in the feeding stage. The sink reactions serve as a nutrient source during the nutrient deprivation or overnight fasting state.

## Metabolic objectives for organs

As a result of the literature search for the organ-specific metabolic pathways, we described each organ by its chief metabolic functions, e.g., Cori cycle between liver and skeletal muscle, arginine synthesis in kidney, citrulline synthesis by the small intestine, cholesterol synthesis by spleen, and vitamin D synthesis by the skin. Glucose from liver enters skeletal muscle, where it is converted to lactate via anaerobic glycolysis. The muscle then releases lactate back into the circulation to be utilized for gluconeogenesis by the liver, contributing to the muscle-liver-Cori cycle (*45*). The kidney is the major organ for the synthesis of arginine from citrulline (*89*). Citrulline synthesized in the small intestine reaches kidney for further metabolism by urea cycle reactions, thereby, contributing to inter-organ amino acid metabolism. Spleen is one of the important hematopoietic organs, and synthesis of dolichol and cholesterol from acetate are important indicators of this process (*90*). The human skin is mainly responsible for the synthesis of vitamin D from 7-dehydrocholesterol in multiple reaction steps (*91*). These physiological functions and their representative biochemical reactions were set as metabolic tasks for each organ (Supplement Table S6).

## Bile composition

Bile salts aid in the digestion and absorption of fat constituents through their micellar properties (*52*). Recon 3D contains the human metabolism of bile acids comprehensively but no organ-specific or circulatory information (*3*). Bile is synthesized in the liver and drained into the gallbladder, via the bile duct. The gallbladder stores the bile constituents, and releases it into the intestinal lumen, i.e., into the duodenum (first part of small intestine) for efficient digestion and absorption of food. To capture bile composition (i.e., presence and absence of exchange metabolites), we followed the large-scale proteomic analysis and specific metabolites measured in human bile (*92, 93*). Conclusive evidence concerning their presence in bile was available for 84 exchange metabolites in Recon 3*; and for 459 exchange metabolites, absence in the bile was concluded (Supplement Table S7). The remaining transport reactions into the bile duct were unconstrained and algorithmically added when extracting the subnetworks depending on their secretion from the gallbladder and its internal metabolism. The storage (aka demand) reactions for 26 bile salts were added to the gallbladder (Supplement Table S7).

## Recon 3* exchange metabolites present/absent in diet

Comprehensive information for the presence in the diet was found for 300 metabolites, and for 50 metabolites the absence in the diet was reported. For the remaining exchange metabolites, no information could be found in the literature. Hence, these were left unconstrained in the meta-reconstruction (Supplement Table S8).

## Metabolomic data

The whole-body metabolic model incorporates 13 extracellular compartments (Table 1). The model was constrained to contain biofluid-specific metabolites for 1105 metabolites that were incorporated into the respective biofluid compartments (i.e., in the blood circulation, portal blood, cerebrospinal fluid, feces, and urine). This was represented by adding the corresponding demand reactions to the biofluids. This information was extracted from various literature references as well as databases, such as HMDB (*19*) (Supplement Table S9). Most of these metabolites were identified as biomarkers in pathological states, and highlight the potential of Harvey in capturing the known biomarkers and prediction of new ones.

## Transport reaction information

Our previous work on human membrane transporters (*94*) served as a compendium of transport proteins. These transport proteins were noted with their organ distribution from the relevant scientific literature. Again, the GPR associations within Recon 3* were used, and the corresponding transport reactions were extracted and incorporated into the core reaction set of the specific organ.

Conclusive evidence for the presence of 166 transport proteins distributed across 26 organs formed the transport protein part of the core reaction set. For the remaining organs, the presence of transport proteins was derived from HPA and HPM. While the presence of the transport protein and its associated reaction was included as core reaction set, the non-occurrence was ignored. This is because the absence of a transport protein across an organ or tissue type is difficult to establish. Interestingly, amino acids transport proteins, ABC transporters, and lipid transporters were found to be more ubiquitously expressed across organs. Most transport protein information was found for kidney, brain, liver, heart, and skeletal muscle, while for hematopoietic cells (e.g., red blood cells, platelets) the least information could be found. We enabled the secretion of mucin degradative products, glycans, and ethanolamine into the large intestinal lumen from the colon (*95, 96*).

## Defining the blood-brain-barrier

We represented the blood-brain-barrier in the WBM reconstructions. The brain is separated from the blood/extracellular compartment by the blood-brain barrier (*97*). This barrier formed by the brain endothelial cells, and exhibit restricted entry of small molecules. Molecules with a molecular mass below 500 Da and possessing high lipid solubility can enter the brain (*98*). This information was used to add the blood-brain-barrier transport reactions for 43 metabolites to the core reaction sets. 240 metabolites have been reported not to pass the blood-brain barrier, which includes lecithin, triglycerides, lysolecithin, and cholesterol (*97, 99*). Thus, the corresponding blood-brain-barrier transport reactions were constrained to zero in the meta-reconstructions, thereby eliminating them from the WBM reconstructions. The remaining transport reactions were unconstrained enabling their addition during the subnetwork generation process. Their addition, therefore, depended on the internal metabolic architecture of the brain (and spinal cord).

## Biomass reactions

The WBM reconstructions contain three different versions of the biomass reaction. These are (i) biomass_reaction, (ii) biomass_maintenance, and (iii) biomass_maintenance_noTrTr. The biomass_reaction is the general biomass reaction as in Recon 3*, biomass_ maintenance is same as biomass_reaction except for the nuclear deoxynucleotides, and biomass_maintenance_noTrTr is devoid of amino acids, nuclear deoxynucleotides, and cellular deoxynucleotides except for adenosine-triphosphate.

The biomass reaction was retained only for tissues known to possess regenerative capacity, i.e., liver (*100*), heart (*101*), and kidney (*102*). For the remaining organs, only biomass_ maintenance was added, indicating the maintenance of cellular metabolic profiles, i.e., the organs capability to synthesize all the biomass components excepting the nuclear deoxynucleotides. The biomass_maintenance_noTrTr reaction was added specifically to model fasting condition. Such a modification was done as the human body has no store for amino acids (*103*). Amino acids if stored intracellularly, increase the osmotic pressure, necessitating their rapid catabolism (*103*). Such catabolic processes mainly occur for those that are not required for protein synthesis.
i. **Published metabolic reconstructions** For the red blood cell (*47*), the adipocytes (*9*), the small intestine (*48*), and the liver (*49*), the reactions present in these published metabolic reconstructions were used to define sets of core reactions. Note that all these reconstructions have been built with Recon 1 (*104*) as a starting point. The published red blood cell reconstruction has been assembled using multiple proteomic data sets (*47*). The published adipocyte reconstruction was generated by tailoring Recon 1 based on genome annotation data, physiological, and biochemical data from online databases (e.g., KEGG (*105*), NCBI, UniProt (*106*), and BRENDA (*107*), and literature (*9*). The liver/hepatocyte reconstruction has been built through manual curation of the relevant scientific literature, using Recon 1 and KEGG as starting points (*49*). Additionally, gene expression datasets of normal human liver samples have served as secondary lines of evidence (*49*). The small intestinal epithelial cell reconstruction (*48*), has been assembled using primary literature, organ-specific books, and databases. Since the small intestinal epithelial cell model maintained different extracellular compartments representing the apical and basolateral polarity of the cell, the reactions were added as such to the core set. However, the GPR association were updated with those used in Recon 3*. Mapping of the reaction content in these published reconstructions onto Recon 3* was done manually using the reaction abbreviation, reaction description, and reaction formula. In the case of the adipocyte, the blood compartment was replaced with the extracellular compartment to find the correct matches in Recon 3* reactions. Additionally, the published adipocyte model (*9*) contained a lumped version of the fatty acid oxidation reactions, hence, the corresponding un-lumped versions were mapped onto Recon 3*. The mapped reactions were added to the core reaction set, after adding the organ-specific prefix to the reaction abbreviations.

### Refinement and curation of the WBM reconstructions

The reaction content of the presented WBM reconstructions have been manually curated at each iteration (more than 100 in total). The algorithmic approach reads in the protein expression per organ and adds the corresponding reaction from Recon 3* to the respective organ. During the building of the whole-body model we encountered genes/proteins that were present in an organ-specific manner as per the human proteome dataset (*12*), but, absent across the respective organs in the draft WBM reconstructions, due to missing transport reactions, which were subsequently added. Note that the development of Recon 3D and the WBM reconstructions occurred in parallel and that Recon 3D contains all those previously missing in Recon 2 (*108*). Therefore, we analyzed the proteins/genes that were not added from the human protein dataset onto the respective organ and included them in the core reaction set per organ. A typical example is the addition of the reactions SALMCOM and SALMCOM2 to the core reaction set for colon, rectum, adrenal gland, platelet, lung, heart, brain, retina, b cells, B-cells, CD4 cells, CD8 cells, NK cells, testis, and prostate. Interestingly, SALMCOM was correctly chosen by the algorithm to be present in liver, gallbladder, pancreas, and kidney. Therefore, we added transport reactions for the participating metabolites of SALMCOM and SALMCOM2, i.e., normete_L and mepi as well as their demand reactions in the blood circulation, provided that both the compounds have been detected in blood and urine (HMDB00819, HMDB04063). This enabled the metabolites to be channeled across the whole body and excreted in the urine by the kidney. Similar to the discussed case, many transport reactions were added during the debugging process that enabled the whole-body routing of phospholipids, cholesterol ester species, and acylcarnitines.

This example shows that the WBM reconstructions can be effectively used to study metabolite cross-talk between organs as well their whole-body physiology. Such intensive manual curation efforts highlight the high quality of the finished models in truly capturing the accurate biochemical picture of the human body and physiologically relevant predictions thereof.

## “Sanity checks” on the WBM reconstructions

Once the reconstruction was converted into a mathematical model following the standards described in (*15*), the model was tested for secretion or production of metabolites when all the exchanges and sinks were shut down. We shall refer to this as leak test. After that, it was checked for the various functions, referred to as function test.

## Leak test

The WBM reconstructions were tested for thermodynamically infeasible loops within the internal reactions that could generate metabolites or compounds when no mass enters the model. Such a test is done in two steps. Firstly, all the exchange, sink, and demand reactions are constrained to zero for the lower bound. Then, all the exchange reactions are optimized to check if the model is carrying any non-zero flux. After that, the biomass is optimized to check if the model is carrying any zero flux. Secondly, a demand reaction for each compartment-specific metabolite in the model is created and optimized. The basic purpose of running such a leak test is to check if the model is generating anything from nothing. In case any of the demand or exchange reactions carry a non-zero flux, the respective reaction is optimized using minNorm (*109*). The flux vector is then analyzed for the reactions that contribute maximally to the defined objective. Such contributing reactions are chosen such that their optimal flux is either equal or greater than the flux of the objective, and their directionality constrained to be irreversible. Typical examples include the production of ATP when no mass enters the cell, and under such conditions, the ATP-utilizing reactions are made irreversible. One should note that this method generates various reactions that can contribute to ATP generation, and only those reactions should be constrained after a through manual inspection. Usually, during the manual reconstruction procedure, a reaction is mentioned as reversible, in case adequate information is unavailable concerning its directionality (*15*).

## Metabolic function tests

During the iterative reconstruction and refinement process, we also tested for the organ-specific metabolic functions (*3*). If no non-zero flux was obtained, we identified the cause and added organ-specific reactions to the draft WBM reconstructions.

## Coupling constraints

Coupling constraints were implemented in Harvey as described previously (*110, 111*). Briefly, coupling constraints enforce that the flux through a set of coupled reactions is proportional to a specified reaction (e.g., biomass). For Harvey, the metabolic and transport reactions in every organ were coupled to the respective organs biomass objective function (BOF). The coupling constraints prevent biologically implausible solutions where the reactions in an organ carry flux even though the flux through the organs BOF is zero. They were realized by implementing a coupling factor of 20000 for each reaction. This allowed each forward and reverse reaction to carry a flux of up to 20000 and −20000 times the flux through the BOF, respectively.

## Simulation details

Please refer to the MATLAB live script for detailed on simulation. MATLAB, Mathworks, Inc.), TheCOBRAToolbox(*109*),tobeobtainedhere: <u>https://opencobra.github.io/cobratoolbox/stable/</u> and the PCSM extension for the COBRA Toolbox.

## Acknowledgments

This study was funded by the Luxembourg National Research Fund (FNR), the ATTRACT programme (FNR/A12/01) and the OPEN programme grants (FNR/O16/11402054), as well as through the National Centre of Excellence in Research (NCER) on Parkinson's disease. The authors are thankful to Dr. N. Poupin, Mrs. C. Clancy, and Mr. M. Ben Guebila, for valuable discussions and editing earlier versions of the manuscript and supplementary material as well as to Dr. E. Schymanski for editing the manuscript. None of the authors have any competing interests.

## Author contributions

IT, RMTF, and SS conceived the study, IT, SS, MKA, and AH contributed to the reconstructions. RMFT, LH, and AN contributed tools and methods. IT performed the simulations. IT, AH, and RMTF analyzed the data. IT, SS, and AH wrote the manuscript. All authors edited the manuscript.

## References

1. D. B. Kell, The virtual human: towards a global systems biology of multiscale, distributed biochemical network models. IUBMB Life 59, 689-695 (2007).

2. P. Hunter et al., A vision and strategy for the virtual physiological human: 2012 update. Interface Focus 3, 20130004 (2013).

3. E. Brunk et al., Recon3D: A Resource Enabling A Three-Dimensional View of Gene Variation in Human Metabolism. Nat Biotech, (Accepted).

4. S. Opdam et al., A Systematic Evaluation of Methods for Tailoring Genome-Scale Metabolic Models. Cell systems 4, 318-329 e316 (2017).

5. S. Magnusdottir et al., Generation of genome-scale metabolic reconstructions for 773 members of the human gut microbiota. Nat Biotechnol 35, 81-89 (2017).

6. H. Kitano, Grand challenges in systems physiology. Frontiers in physiology 1, 3 (2010).

7. B. Palsson, Systems biology: properties of reconstructed networks. (Cambridge University Press, Cambridge, 2006), pp. xii, 322 p.

8. J. D. Orth, I. Thiele, B. O. Palsson, What is flux balance analysis? Nat Biotechnol 28, 245-248 (2010).

9. A. Bordbar et al., A multi-tissue type genome-scale metabolic network for analysis of whole-body systems physiology. BMC systems biology 5, 180 (2011).

10. A. Nilsson, A. Mardinoglu, J. Nielsen, Predicting growth of the healthy infant using a genome scale metabolic model. NPJ Syst Biol Appl 3, 3 (2017).

11. A. Heinken, I. Thiele, Systematic prediction of health-relevant human-microbial co-metabolism through a computational framework. Gut Microbes 6, 120-130 (2015).

12. M. S. Kim et al., A draft map of the human proteome. Nature 509, 575-581 (2014).

13. M. Uhlen et al., Proteomics. Tissue-based map of the human proteome. Science (New York, N.Y) 347, 1260419 (2015).

14. N. Vlassis, M. P. Pacheco, T. Sauter, Fast reconstruction of compact context-specific metabolic network models. PLoS Comput Biol 10, e1003424 (2014).

15. I. Thiele, B. O. Palsson, A protocol for generating a high-quality genome-scale metabolic reconstruction. Nature protocols 5, 93-121 (2010).

16. W. S. Snyder et al., Report on the Task Group on Reference Man. INTERNATIONAL COMMISSION ON RADIOLOGICAL PROTECTION No 23 (1975).

17. A. Noronha et al., ReconMap: an interactive visualization of human metabolism. Bioinformatics (Oxford, England), (2016).

18. M. K. Aurich, I. Thiele, Computational Modeling of Human Metabolism and Its Application to Systems Biomedicine. Methods in molecular biology (Clifton, N.J 1386, 253-281 (2016).

19. D. S. Wishart et al., HMDB 3.0‐‐The Human Metabolome Database in 2013. Nucleic Acids Res 41, D801-807 (2013).

20. I. Elmadfa, Österreichischer Ernährungsbericht 2012. (Vienna, ed. 1., 2012).

21. R. Milo, P. Jorgensen, U. Moran, G. Weber, M. Springer, BioNumbers‐‐the database of key numbers in molecular and cell biology. Nucleic Acids Res 38, D750-753 (2010).

22. T. Skotland et al., Determining the Turnover of Glycosphingolipid Species by Stable-Isotope Tracer Lipidomics. J Mol Biol 428, 4856-4866 (2016).

23. H. Wackerhage et al., Recovery of free ADP, Pi, and free energy of ATP hydrolysis in human skeletal muscle. Journal of applied physiology 85, 2140-2145 (1998).

24. M. D. Mifflin et al., A new predictive equation for resting energy expenditure in healthy individuals. Am J Clin Nutr 51, 241-247 (1990).

25. J. M. Berg, J. L. Tymoczko, L. Stryer, Biochemistry. (W H Freeman, New York, ed. 5th 2002).

26. H. K. Biesalski, P. Grimm, Pocket Atlas of Nutrition. (Thieme, 2005).

27. J. P. DeLany, D. E. Kelley, K. C. Hames, J. M. Jakicic, B. H. Goodpaster, High energy expenditure masks low physical activity in obesity. International journal of obesity 37, 1006-1011 (2013).

28. C. Tudor-Locke, Y. Hatano, R. P. Pangrazi, M. Kang, Revisiting “how many steps are enough?”. Medicine and science in sports and exercise 40, S537-543 (2008).

29. M. A. van Baak, Physical activity and energy balance. Public health nutrition 2, 335-339 (1999).

30. C. Koutsari, M. S. Mundi, A. H. Ali, M. D. Jensen, Storage rates of circulating free fatty acid into adipose tissue during eating or walking in humans. Diabetes 61, 329-338 (2012).

31. G. Clarke et al., Minireview: Gut microbiota: the neglected endocrine organ. Mol Endocrinol 28, 1221-1238 (2014).

32. T. H. M. P. Consortium, A framework for human microbiome research. Nature 486, 215-221 (2012).

33. A. Heinken et al., Personalized modeling of the human gut microbiome reveals distinct bile acid deconjugation and biotransformation potential in healthy and IBD individuals. BioRxiv preprint, (2017).

34. P. Spanogiannopoulos, E. N. Bess, R. N. Carmody, P. J. Turnbaugh, The microbial pharmacists within us: a metagenomic view of xenobiotic metabolism. Nature reviews 14, 273-287 (2016).

35. H. Li et al., The outer mucus layer hosts a distinct intestinal microbial niche. Nat Commun 6, 8292 (2015).

36. M. G. Neuman et al., Alcohol, microbiome, life style influence alcohol and non-alcoholic organ damage. Exp Mol Pathol 102, 162-180 (2017).

37. P. Louis, H. J. Flint, Diversity, metabolism and microbial ecology of butyrate-producing bacteria from the human large intestine. FEMS microbiology letters 294, 1-8 (2009).

38. D. A. Beard et al., Multiscale modeling and data integration in the virtual physiological rat project. Ann Biomed Eng 40, 2365-2378 (2012).

39. M. Viceconti et al., “in silico Clinical Trials: How Computer Simulation will Transform the Biomedical Industry,” (2016).

40. J. R. Karr et al., A whole-cell computational model predicts phenotype from genotype. Cell 150, 389-401 (2012).

41. M. Krauss et al., Integrating cellular metabolism into a multiscale whole-body model. PLoS Comput Biol 8, e1002750 (2012).

42. J. L. Sonnenburg, F. Backhed, Diet-microbiota interactions as moderators of human metabolism. Nature 535, 56-64 (2016).

43. S. Shoaie et al., Quantifying Diet-Induced Metabolic Changes of the Human Gut Microbiome. Cell metabolism 22, 320-331 (2015).

44. S. P. Claus et al., Systemic multicompartmental effects of the gut microbiome on mouse metabolic phenotypes. Molecular systems biology 4, 219 (2008).

45. R. K. Murray, D. K. Granner, P. A. Mayes, V. W. Rodwell, A Lange Medical book: Harper's Biochemistry. (Appleton and Lange, ed. 25th, 2000), pp. 298-305.

46. S. A. S. Gropper, J. L. Smith, J. L. Groff, Advanced nutrition and human metabolism. (Wadsworth/Cengage Learning, Australia; United States, ed. 5th ed., 2009).

47. A. Bordbar, N. Jamshidi, B. O. Palsson, iAB-RBC-283: A proteomically derived knowledge-base of erythrocyte metabolism that can be used to simulate its physiological and patho-physiological states. BMC systems biology 5, 110 (2011).

48. S. Sahoo, I. Thiele, Predicting the impact of diet and enzymopathies on human small intestinal epithelial cells. Hum Mol Genet 22, 2705-2722 (2013).

49. C. Gille et al., HepatoNet1: a comprehensive metabolic reconstruction of the human hepatocyte for the analysis of liver physiology. Molecular systems biology 6, 411 (2010).

50. M. Uhlen et al., Towards a knowledge-based Human Protein Atlas. Nat Biotechnol 28, 1248-1250 (2010).

51. P. Flicek et al., Ensembl 2014. Nucleic Acids Res 42, D749-755 (2014).

52. R. K. Murray et al., Harper's illustrated Biochemistry. (Mc Graw Hill, ed. 28th, 2009), pp. 346-351.

53. N. B. Ruderman, Muscle amino acid metabolism and gluconeogenesis. Annu Rev Med 26, 245-258 (1975).

54. N. J. Cano, D. Fouque, X. M. Leverve, Application of branched-chain amino acids in human pathological states: renal failure. J Nutr 136, 299s-307s (2006).

55. H. Nieth, P. Schollmeyer, Substrate-utilization of the human kidney. Nature 209, 1244-1245 (1966).

56. M. Gold, J. J. Spitzer, Metabolism of free fatty acids by myocardium and kidney. Am J Physiol 206, 153-158 (1964).

57. C. P. Berg et al., Human mature red blood cells express caspase-3 and caspase-8, but are devoid of mitochondrial regulators of apoptosis. Cell death and differentiation 8, 1197-1206 (2001).

58. P. Delva, M. Degan, M. Trettene, A. Lechi, Insulin and glucose mediate opposite intracellular ionized magnesium variations in human lymphocytes. J Endocrinol 190, 711-718 (2006).

59. W. Jia, H. H. Pua, Q. J. Li, Y. W. He, Autophagy regulates endoplasmic reticulum homeostasis and calcium mobilization in T lymphocytes. J Immunol 186, 1564-1574 (2011).

60. W. Jia, Y. W. He, Temporal regulation of intracellular organelle homeostasis in T lymphocytes by autophagy. J Immunol 186, 5313-5322 (2011).

61. K. Kaneda, M. Kataoka, T. Kishiye, H. Yamamoto, K. Wake, The intracellular distribution of cell organelles in natural killer cells during the cytolysis of bound tumor cells, with special reference to the rod-cored vesicles. Arch Histol Cytol 54, 69-79 (1991).

62. E. C. Dell'Angelica, C. Mullins, S. Caplan, J. S. Bonifacino, Lysosome-related organelles. FASEB J 14, 1265-1278 (2000).

63. P. Roda-Navarro, H. T. Reyburn, The traffic of the NKG2D/Dap10 receptor complex during natural killer (NK) cell activation. J Biol Chem 284, 16463-16472 (2009).

64. A. J. Marcus, The role of lipids in platelet function: with particular reference to the arachidonic acid pathway. J Lipid Res 19, 793-826 (1978).

65. A. J. Marcus, D. Zucker-Franklin, L. B. Safier, H. L. Ullman, Studies on human platelet granules and membranes. J Clin Invest 45, 14-28 (1966).

66. J. Polasek, Platelet lysosomal acid phosphatase enzyme activity as a marker of platelet procoagulant activity. Blood Transfus 7, 155-156 (2009).

67. S. Zharikov, S. Shiva, Platelet mitochondrial function: from regulation of thrombosis to biomarker of disease. Biochem Soc Trans 41, 118-123 (2013).

68. R. I. Handin, S. E. Lux, T. P. Stossel, Blood: Principles and Practice of Hematology. (Lippincot Williams & Wilkins, USA, ed. 2nd, 2003).

69. K. M. Meyers, C. D. Barnes, The platelet amine storage granule. (CRC Press, Inc, Florida, USA, 1992).

70. T. Takahashi et al., Vitrification of human monocytes. Cryobiology 23, 103-115 (1986).

71. F. W. van der Meulen, M. Reiss, E. A. Stricker, E. van Elven, A. E. von dem Borne, Cryopreservation of human monocytes. Cryobiology 18, 337-343 (1981).

72. S. Sekine, M. Miura, T. Chihara, Organelles in developing neurons: essential regulators of neuronal morphogenesis and function. Int J Dev Biol 53, 19-27 (2009).

73. S. Weiss et al., Multipotent CNS stem cells are present in the adult mammalian spinal cord and ventricular neuroaxis. J Neurosci 16, 7599-7609 (1996).

74. Y. Imanishi, V. Gerke, K. Palczewski, Retinosomes: new insights into intracellular managing of hydrophobic substances in lipid bodies. J Cell Biol 166, 447-453 (2004).

75. A. Kaasik et al., Energetic crosstalk between organelles: architectural integration of energy production and utilization. Circ Res 89, 153-159 (2001).

76. J. Piquereau et al., Mitochondrial dynamics in the adult cardiomyocytes: which roles for a highly specialized cell? Frontiers in physiology 4, 102 (2013).

77. M. Malatesta et al., Ultrastructural morphometrical and immunocytochemical analyses of hepatocyte nuclei from mice fed on genetically modified soybean. Cell Struct Funct 27, 173-180 (2002).

78. S. A. Nottola et al., Cryopreservation and xenotransplantation of human ovarian tissue: an ultrastructural study. Fertil Steril 90, 23-32 (2008).

79. K. Hoffmann et al., New application of a subcellular fractionation method to kidney and testis for the determination of conjugated linoleic acid in selected cell organelles of healthy and cancerous human tissues. Anal Bioanal Chem 381, 1138-1144 (2005).

80. S. Karnati, E. Baumgart-Vogt, Peroxisomes in airway epithelia and future prospects of these organelles for pulmonary cell biology. Histochem Cell Biol 131, 447-454 (2009).

81. D. P. Boshier, H. Holloway, Morphometric analyses of adrenal gland growth in fetal and neonatal sheep. III. Volumes of the major organelles within zona fasciculata steroidogenic cells. J Anat 178, 175-187 (1991).

82. K. Yoshida, K. Katayanagi, Y. Kawamura, K. Saito, Y. Nakanuma, Reestablishment of rabbit gallbladder epithelial cells in collagen gel culture and their alterations by cytochalasin B and transforming growth factor beta-1. A morphologic study. Pathol Res Pract 192, 634-645 (1996).

83. T. J. Li et al., Basaloid squamous cell carcinoma of the esophagus with or without adenoid cystic features. Arch Pathol Lab Med 128, 1124-1130 (2004).

84. F. E. Pittman, J. C. Pittman, An electron microscopic study of epithelium of normal human sigmoid colonic mucosa. Gut 7, 644-661 (1966).

85. L. Napolitano, The Differentiation of White Adipose Cells. An Electron Microscope Study. J Cell Biol 18, 663-679 (1963).

86. P. M. Novikoff, A. B. Novikoff, Peroxisomes in absorptive cells of mammalian small intestine. J Cell Biol 53, 532-560 (1972).

87. L. K. Summers et al., Uptake of individual fatty acids into adipose tissue in relation to their presence in the diet. Am J Clin Nutr 71, 1470-1477 (2000).

88. S. A. Lanham-New, I. A. MacDonald, E. M. Roche, Nutrition and Metabolism. (Wiley-Blackwell, ed. 2nd, 2010).

89. M. C. van de Poll, P. B. Soeters, N. E. Deutz, K. C. Fearon, C. H. Dejong, Renal metabolism of amino acids: its role in interorgan amino acid exchange. Am J Clin Nutr 79, 185-197 (2004).

90. J. E. Potter, M. J. James, A. A. Kandutsch, Sequential cycles of cholesterol and dolichol synthesis in mouse spleens during phenylhydrazine-induced erythropoiesis. J Biol Chem 256, 2371-2376 (1981).

91. T. C. Chen et al., Factors that influence the cutaneous synthesis and dietary sources of vitamin D. Arch Biochem Biophys 460, 213-217 (2007).

92. F. Fuda, S. B. Narayan, R. H. Squires,Jr., M. J. Bennett, Bile acylcarnitine profiles in pediatric liver disease do not interfere with the diagnosis of long-chain fatty acid oxidation defects. Clin Chim Acta 367, 185-188 (2006).

93. A. Farina, J. M. Dumonceau, P. Lescuyer, Proteomic analysis of human bile and potential applications for cancer diagnosis. Expert Rev Proteomics 6, 285-301 (2009).

94. S. Sahoo, M. K. Aurich, J. J. Jonsson, I. Thiele, Membrane transporters in a human genome-scale metabolic knowledgebase and their implications for disease. Frontiers in physiology 5, 91 (2014).

95. D. E. Chang et al., Carbon nutrition of Escherichia coli in the mouse intestine. Proceedings of the National Academy of Sciences of the United States of America 101, 7427-7432 (2004).

96. A. J. Fabich et al., Comparison of carbon nutrition for pathogenic and commensal Escherichia coli strains in the mouse intestine. Infect Immun 76, 1143-1152 (2008).

97. Z. Redzic, Molecular biology of the blood-brain and the blood-cerebrospinal fluid barriers: similarities and differences. Fluids Barriers CNS 8, 3 (2011).

98. W. M. Pardridge, The blood-brain barrier: bottleneck in brain drug development. NeuroRx 2, 3-14 (2005).

99. W. M. Pardridge, L. J. Mietus, Palmitate and cholesterol transport through the blood-brain barrier. J Neurochem 34, 463-466 (1980).

100. H. Malhi, A. N. Irani, S. Gagandeep, S. Gupta, Isolation of human progenitor liver epithelial cells with extensive replication capacity and differentiation into mature hepatocytes. J Cell Sci 115, 2679-2688 (2002).

101. S. E. Senyo et al., Mammalian heart renewal by pre-existing cardiomyocytes. Nature 493, 433-436 (2013).

102. Y. Li, R. A. Wingert, Regenerative medicine for the kidney: stem cell prospects & challenges. Clinical and translational medicine 2, 11 (2013).

103. S. Lanham-New, I. MacDonald, H. Roche, Nutrition and metabolism. (Wiley, ed. 2nd, 2013).

104. N. C. Duarte et al., Global reconstruction of the human metabolic network based on genomic and bibliomic data. Proceedings of the National Academy of Sciences of the United States of America 104, 1777-1782 (2007).

105. S. Okuda et al., KEGG Atlas mapping for global analysis of metabolic pathways. Nucleic Acids Res, (2008).

106. T. U. Consortium, Reorganizing the protein space at the Universal Protein Resource (UniProt). Nucleic Acids Res 40, D71-75 (2012).

107. M. Scheer et al., BRENDA, the enzyme information system in 2011. Nucleic Acids Res 39, D670-676 (2011).

108. I. Thiele et al., A community-driven global reconstruction of human metabolism. Nat Biotechnol 31, 419-425 (2013).

109. L. Heirendt et al., Creation and analysis of biochemical constraint-based models: the COBRA Toolbox v3.0. arXiv preprint, (2017).

110. A. Heinken, S. Sahoo, R. M. Fleming, I. Thiele, Systems-level characterization of a host-microbe metabolic symbiosis in the mammalian gut. Gut Microbes 4, 28-40 (2013).

111. I. Thiele, R. M. Fleming, A. Bordbar, J. Schellenberger, B. O. Palsson, Functional characterization of alternate optimal solutions of Escherichia coli's transcriptional and translational machinery. Biophysical journal 98, 2072-2081 (2010).

